# Multiscalar electrical spiking in *Schizophyllum commune*

**DOI:** 10.1101/2023.04.20.537646

**Authors:** Andrew Adamatzky, Ella Schunselaar, Han A. B. Wösten, Phil Ayres

## Abstract

Growing colonies of the split-gill fungus *Schizophyllum commune* show action potential-like spikes of extracellular electrical potential. We analysed several days of electrical activity recording of the fungus and discovered three families of oscillatory patterns. Very slow activity at a scale of hours, slow activity at a scale of tens minutes and very fast activity at scale of half-minute. We simulated the spiking behaviour using FitzHugh-Nagume model, uncovered mechanisms of spike shaping. We speculated that spikes of electrical potential might be associated with transportation of nutrients and metabolites.

## 1. Introduction

Organisms generate electromagnetic fields and employ the fields to obtain and fuse information about their environment, to establish communication between components of their bodies and to control the shape of their bodies [1, 2, 3, 4, 5, 6, 7, 8, 9, 10]. From an information processing point of view, one from the most interesting phenomena of bio-electricity is neural spiking. Spikes of electrical potential are the most well-known attributes of neurons and are attributed to their learning and decision making [11, 12, 13]. Not only neurons, but most living substrates can produce spikes of electrical potential. These include Protozoa [14, 15, 16], Hyrdoroza [17], slime moulds [18, 19] and plants [20, 21, 22]. Action potential-like spiking activity in fungi was first documented in 1976 [23], further confirmed in 1995 [24] and techniques for recording the electrical activity in fruiting bodies and colonised substrates was identified in 2018 [25]. While trying to uncover mechanisms of integrative electrical communication in fungi, we recorded and analysed electrical activity of oyster fungi *Pleurotus djamor* [25], bracket fungi *Ganoderma resinaceum* [26], ghost fungi (*Omphalotus nidiformis*), Enoki fungi (*Flammulina velutipes*), split gill fungi (*Schizophyllum commune*) and caterpillar fungi (*Cordyceps militaris*) [27]. We found significant degrees of variability of electrical spiking characteristics and substantial complexity of the electrical communication events [28]. In the experiments mentioned above we inserted electrodes in a substrate colonised by fungi. Topologies of the mycelium networks inside the colonised substrates have not been disclosed which complicated the interpretation of spiking activity. Therefore, we decided to conduct experiments on more homogeneous fungal material: colonies cultured on agar in Petri dishes. Results of this study are reported below.

## 2. Methods

Split-gill fungus *Schizophyllum commune*, strain H4-8A (Utrecht University, The Netherlands) was grown on *S. commune* minimal medium (SCMM) [29] supplemented with 1.5% agar for three days at 30^*°*^C in the dark. Petri dishes with fungal colonies were not opened during experiments. We melted openings in the Petri dish lids with a hot needle and inserted electrodes until they touched the bottom of the dishes 1a. Electrical activity of the fungal colonies was recorded using pairs of iridium-coated stainless steel sub-dermal needle electrodes (Spes Medica S.r.l., Italy), with twisted cables and ADC-24 (Pico Technology, UK) high-resolution data logger with a 24-bit A/D converter. Galvanic isolation and software-selectable sample rates all contribute to a superior noise-free resolution. Each pair of electrodes reported a potential difference between the electrodes. In each pair of differential electrodes, the distance between electrodes was c. 10 mm. We recorded electrical activity at one sample per second. During the recording, the logger has been doing as many measurements as possible (typically up to 600 per second) and saving the average value. The acquisition voltage range was 78 mV. The recording continued for nearly 6 days. We have collected data from 16 pairs of differential electrodes. Due to inert coating the electrodes did not interfere with growth of the colonies as illustrated in Fig. 1b. Spiking activity of very slow and slow spikes (see definitions below) has be analysed manually due to low number of spikes. Fast spikes of electrical potential have been detected in a semi-automatic mode as follows; for each sample measurement *x*_*i*_ we calculated the average value of its neighbourhood as 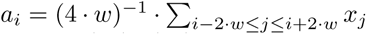. The index *i* is considered a peak of the local spike if |*x*_*i*_| *−* |*a*_*i*_| *> δ*. The list of spikes were further filtered by removing false spikes located at a distance *d* from a given spike. Parameters specific to *S. commune* are *w* = 20, *δ* = 0.01, *d* = 30. An example of spike detection is shown in Fig. 2. The detection is not 100% (as any other algorithms) but over 90% having been detected, as per manual checks.

**Figure 1:**
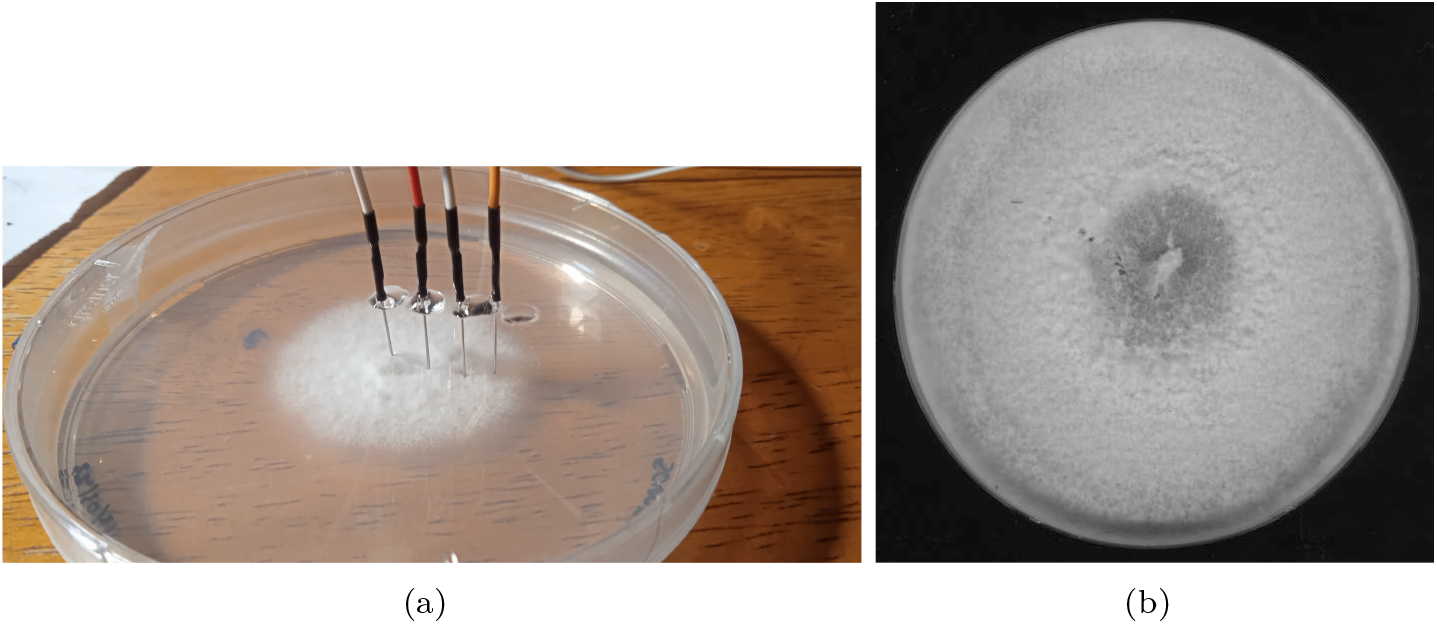
Experimental setup of recording electrical activity of the colony of *S. commune*. (a) Pairs of electrodes are positioned in contact with agar gel through the openings in the Petri dish’s lid and fixed with a glue. (b) Scan of the colony after ten days of the recording.

**Figure 2:**
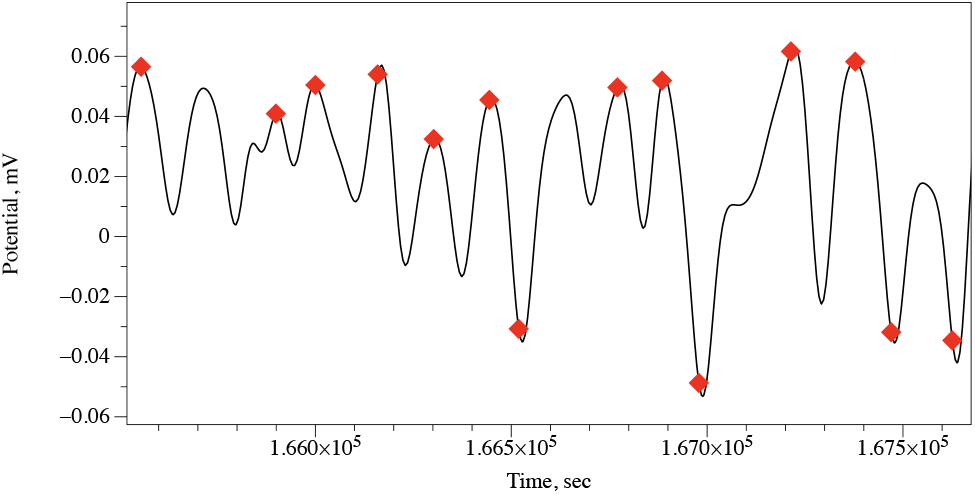
Example of spike detection.

To illustrate a nature of electrical potential spikes we used FitzHugh-Nagumo (FHN) equations [30, 31, 32]. FHN model is which are a qualitative approximation of the Hodgkin-Huxley model [33] of electrical activity of living cells:

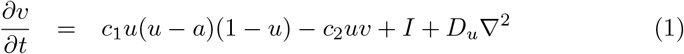

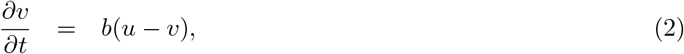

where *u* is a value of a trans-membrane potential, *v* a variable accountable for a total slow ionic current, or a recovery variable responsible for a slow negative feedback, *I* is a value of an external stimulation current. We integrated the system using the Euler method with the five-node Laplace operator, a time step Δ*t* = 0.015 and a grid point spacing Δ*x* = 2, while other parameters were *D*_*u*_ = 1, *a* = 0.13, *b* = 0.013, *c*_1_ = 0.26. We controlled excitability of the medium by varying *c*_2_ from 0.05 (fully excitable) to 0.015 (non excitable). Boundaries are considered to be impermeable: *∂u/∂***n** = 0, where **n** is a vector normal to the boundary. We simulated electrodes by calculating a potential 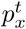 at an electrode location as 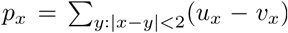. The simulation was conducted on a grid 300*×*300 nodes. Source of the one-off excitation was placed in the middle of the grid, coordinates (150,150). Active electrode *x* was placed at coordinates (190,150) and reference electrode *y* at coordinates (190+d,150), where *d* = 1, 5, 10, 40, 80.

## 3. Results

Typically, recordings obtained in experiments consist for 80% of duration of irregular patterns of wide band electrical activity with no pronounced regular patterns. However, there are segments of activity, where trains of regular spikes of electrical potential are presented. This is illustrated in experimental recording plotted in Fig. 3, where a train of spikes is shown in the magnified insert.

**Figure 3:**
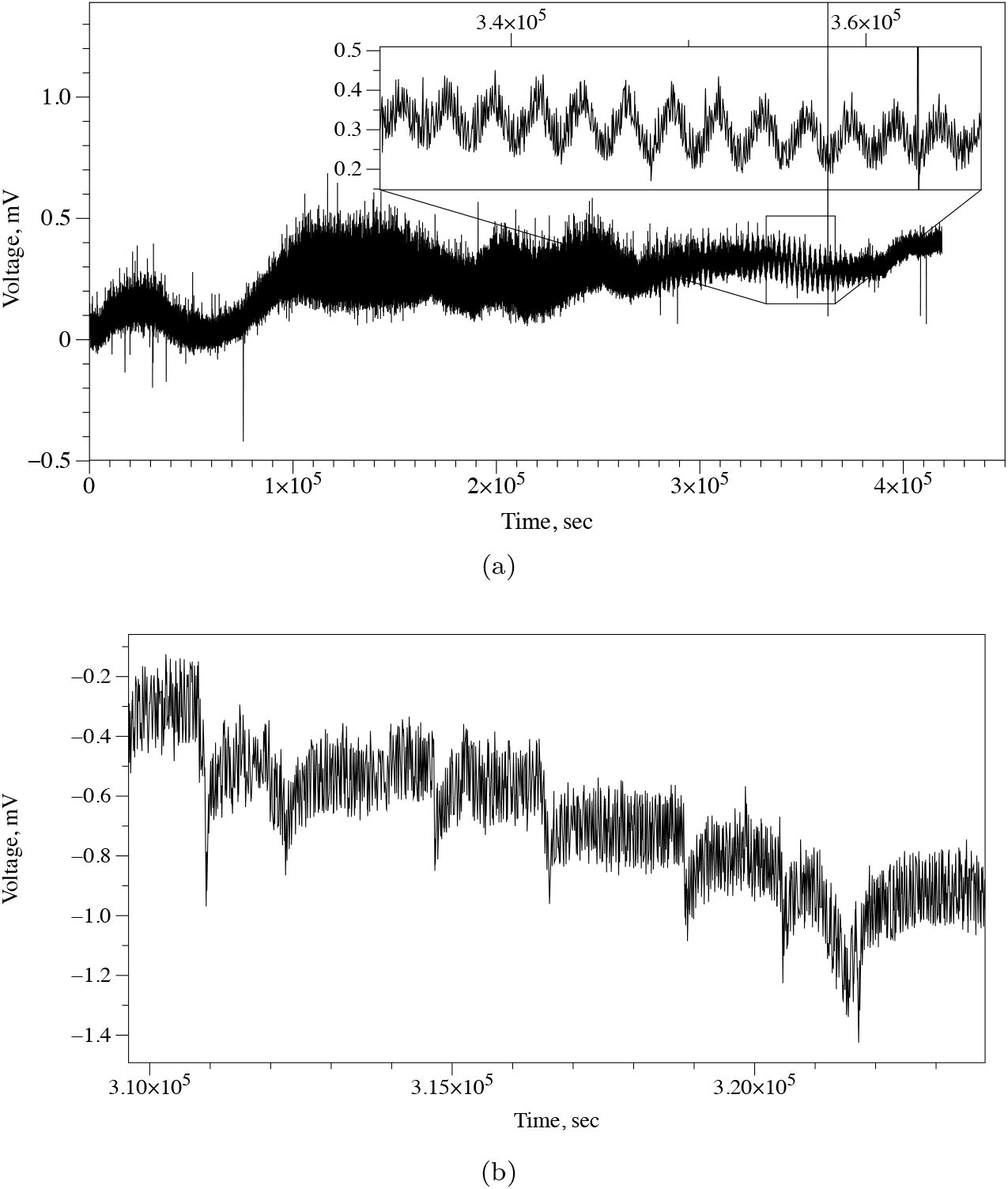
Emergence of regular electrical activity in cultures of *S. commune*. (a) Train of symmetrical spikes is magnified in the insert. (b) Action potential-like spikes.

Manual analysis of the recordings established three scales of electrical potential oscillations: (1) very slow spikes, an hour range, (2) fast spikes, few minutes range, and (3) very fast spikes, less than half a minute range.

### Very slow spikes

In trains of symmetrical spikes, an average duration of a spike is 2573 sec (median 2630 sec, *σ*=168) and an average amplitude is 0.16 mV (median 0.16 mV, *σ*=0.02). An average distance between spikes is 2656 sec (median 2630 sec, *σ*=278). Each spike is symmetrical, i.e. a temporal distance from the start of the spike to the summit is equal to a temporal distance from the summit to the end of the spike.

### Slow spikes

A train of action potential-like spikes is illustrated in Fig. 3b. An average duration of a spike is 457 sec (median 424 sec, *σ*=120), average amplitude is 0.4 mV (median 0.4 mV, *σ*=0.10). An average distance between spikes is 1819 sec (median 1738 sec, *σ*=532). The spikes are temporally asymmetric: average duration from start of a spike to its summit is 89 sec (media 83 sec, *σ*=59). That is 20% of the average width of a spike.

### Very fast spikes

Third family of action potential-like spikes discovered are fast spikes (Fig. 4a). Average width of fast spikes is 24 sec (median 23 sec, *σ*=0.07). Average amplitude is 0.36 mV (media 0.35 sec, *σ*=0.06). In the fragment illustrated in Fig. 4a, an average distance between spikes is 148 sec (media 143 sec, *σ*=68). However, in the overall recording of 5 days (Fig. 4b), the characteristics of the spikes vary substantially. A bar-code representation of the spikes is shown in Fig. 4c. Distance between spikes varies from 65 sec to 114 min (Fig. 4d). The spikes do appear in trains, an average train length is six spikes and median four spikes, *σ*=8.1.

**Figure 4:**
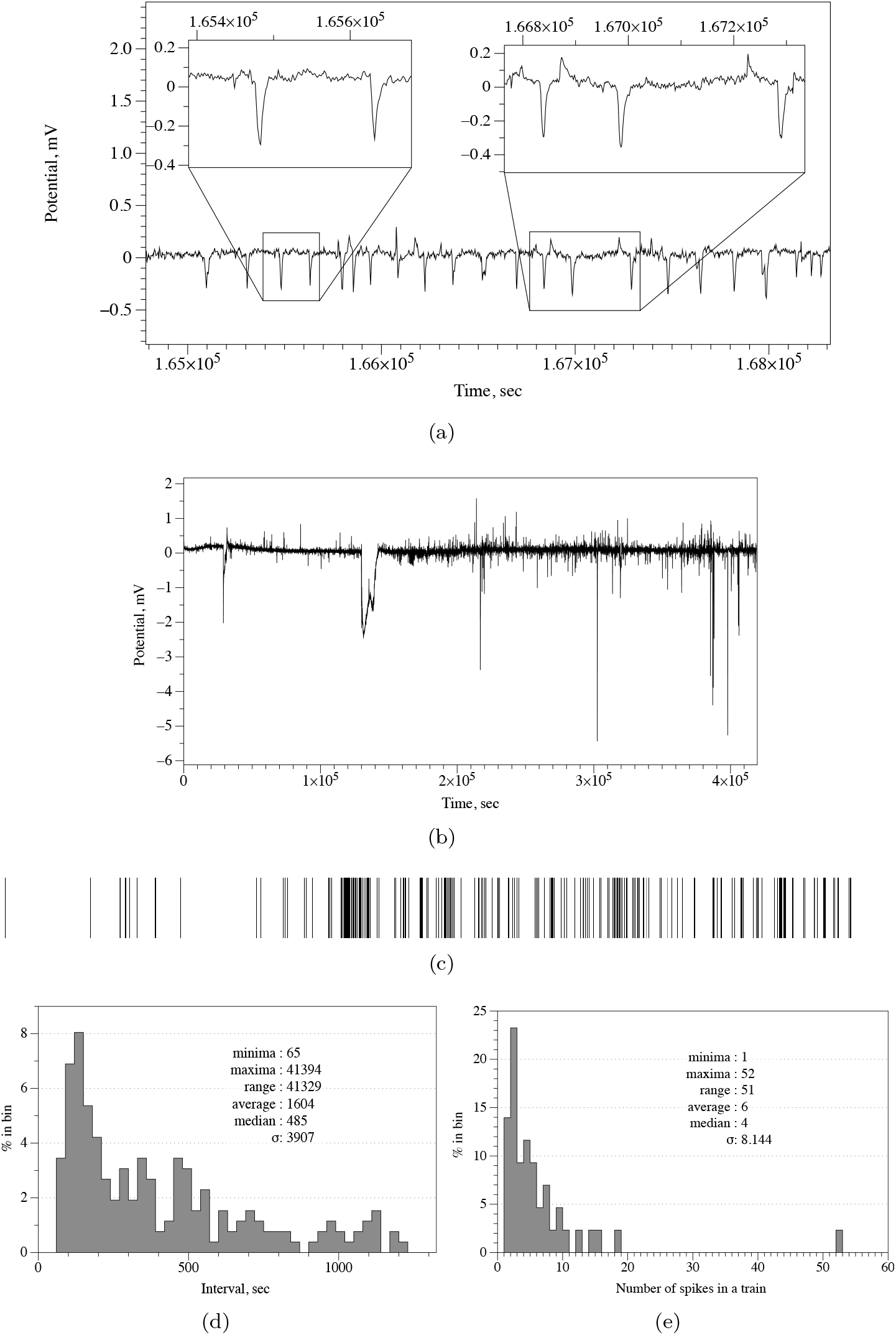
Fast action potential-like spikes in cultures of *S. commune*. (a) Example of a train of spikes with most characteristic spikes zoomed in the insert. (b) Recording conducted during five days. (c) Spikes detected represented as bar code. (b) Distribution of distances between fast spikes.

To demonstrate potential mechanisms of spiking we used Fitz-HughNagumo (FHN) model. We represented a source of electrical excitation at the centre of the colony. Due to the colony being nearly homogeneous — at the macro scale – the wave of excitation propagates as a circular wave (Fig. 5a). When a wave of excitation crosses a loci between the electrodes, a potential difference is recorded. This is illustrated in Fig. 5. In this example, the left electrode is active and the right electrode is a reference. A potential is zero when an excitation wave is far from the electrodes (Fig. 5b). As soon as the wave front starts propagating under the active electrode a positive electrical potential is recorded (Fig. 5cd). When the wave front approaches the reference electrodes, the value of the electrical potential recorded on the active electrode drops to below zero (Fig. 5ef). When a negatively charged tail of the wave front passes under the reference electrode, a second peak of positive electrical potential is recorded on the active electrode (Fig. 5gh). In Fig. 6, we illustrate how the distance *d* between electrodes might affect the shape of the spike recorded. A spike has two extremes when *d* = 1, three extremes for *d* = 5, 10, and five extremes when *d* = 40, 80. Indeed, we should keep in mind that an excitation front width in the model was c. 15 nodes.

**Figure 5:**
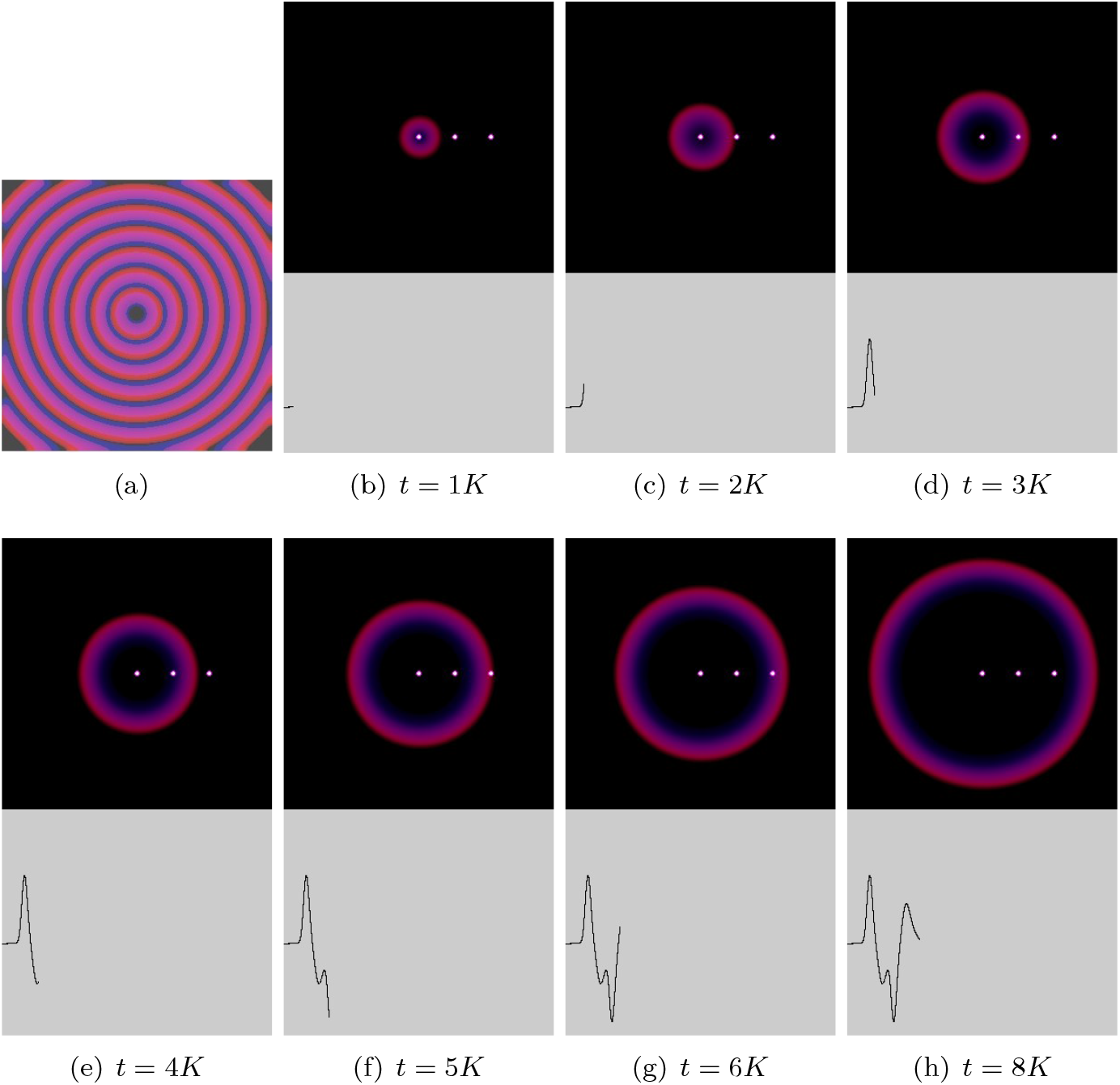
Modelling spiking activity. (a) Propagation of excitation wave in a conducive space imitating colony of *S. commune*. The image is an overlap of time lapse snapshots of a single wave, saved every 1500th iteration. (b–j) Snapshots of excitation dynamics recorded at 1K iterations of numerical integration of FHN model to 9K iterations. Left white dot is a source of excitation. Two right white dots is a pair of differential electrodes. A potential difference between the electrodes is shown at the bottom of each snapshot. Distance between electrodes is 40 nodes of the integration grid.

**Figure 6:**
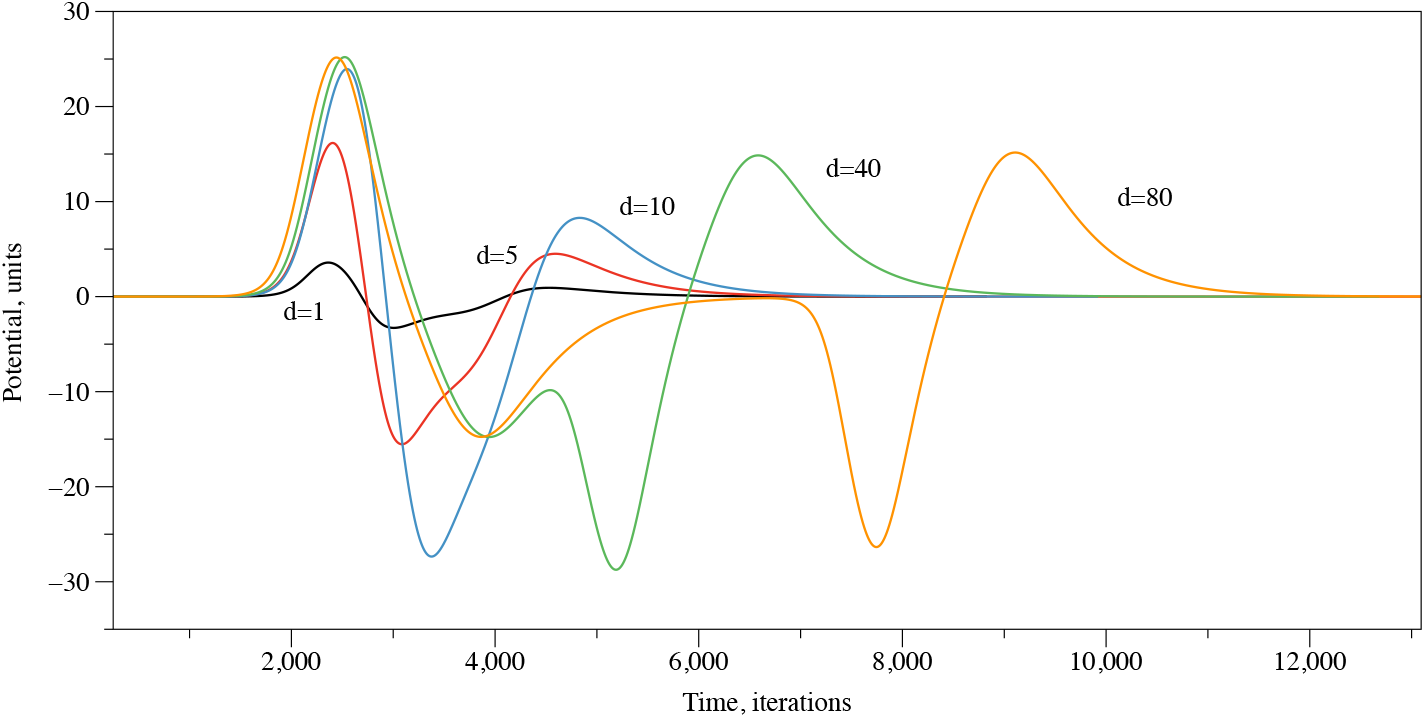
Spikes recorded in simulation with various distance between electrodes.

## 4. Discussion

We observed three families of regular electrical activity in colonies of *S. commune*. We observed very slow spikes with a duration of c. 43 min, slow spikes with a duration of c. 8 min and very fast spikes of a c. 24 sec duration, on average. Thus, the three temporal scales of spiking represent three types of waves, or propagating patterns, which causes changes in the electrical potential. What could be a reason for such a high degree of variability in the speed of propagating excitation? A translocation of metabolites could be the most realistic answer. A study on continuous imaging of amino-acid translocation in mycelia of *Phanerochaete velutina* [34] uncovered pulsating fluxes in which the speed of propagation varies from 9 to 51 mm/h. In our experiments, a distance between electrodes in each differential pair was about 10 mm. This means that c. 9 mm/h translocation speed, reported in [34], matches the very slow spikes of electrical potential that were measured. The highest speed of translocation, 51 mm/h, roughly matches the slow spikes with an average duration of 8 min. Moreover, slow spikes with 8 min duration can be also be related to propagation of calcium waves. Assuming the fastest calcium wave, as per review [35], propagates at 0.03 mm/sec, the wave will travel between electrodes in c. 5 mins. Very fast spikes, indicating a speed of excitation propagation of c. 1500 mm/h are unlikely to be related to transport of metabolites. However, there is a sufficient amount of experimental evidence, see an overview in [36, 37], that fungi grow in a pulsating manner rather than at a constant rate. Namely, *Neurospora crassa* shows 3-6 sec pulse, and a speed of 201 nm/sec, while *Trichoderma virde* 4-6 sec pulse and 201 nm/s speed [36]. In seven species of fungi studied in [36], pulsing varies from 4 to 30 sec in duration. We can therefore hypothesise that fast action potential spiking is associated with, or even controls, a pulsating growth of *S. commune*.

The wave of electrical activity propagating in the fungal colonies, might also be employed in sensorial fusion, information transfer, and distributed decision making. We will explore this possibility in our future research.

## 5. Acknowledgement

The research has been conducted under the framework of the FUNGATERIA (www.fungateria.eu) project, which has received funding from the European Union’s HORIZON-EIC-2021-PATHFINDER CHALLENGES programme under grant agreement No. 101071145. It is co-funded by the UK Research and Innovation grant No. 10048406.

## References

[1] R. Beutner, Source of bioelectricity, investigated by the relation between stainability and electric charges in tissues and artificial models, Proceedings of the Society for Experimental Biology and Medicine 27 (1) (1929) 44–46.

[2] K. S. Cole, H. J. Curtis, Bioelectricity: electric physiology, Medical physics 2 (1950) 82–90.

[3] H. Burr, Bioelectricity: potential gradients, Medical physics 2 (1950) 90–94.

[4] R. Beutner, J. Lozner, The relation of life to electricity: Part iii stainability and electromotive forces of protein; the influence of watersoluble acids, Protoplasma 12 (1931) 145–160.

[5] W. Dorfman, Electrical polarity of the amphibian egg and its reversal through fertilization, Protoplasma 21 (1934) 245–257.

[6] R. Beutner, J. Lozner, The relation of life to electricity: Part viii the mechanism of oxidation-reduction potentials in living tissues, Protoplasma 19 (1933) 370–380.

[7] J. L. Whited, M. Levin, Bioelectrical controls of morphogenesis: from ancient mechanisms of cell coordination to biomedical opportunities, Current opinion in genetics & development 57 (2019) 61–69.

[8] M. Levin, Bioelectric mechanisms in regeneration: unique aspects and future perspectives, in: Seminars in cell & developmental biology, Vol. 20, Elsevier, 2009, pp. 543–556.

[9] M. Levin, Bioelectric signaling: Reprogrammable circuits underlying embryogenesis, regeneration, and cancer, Cell 184 (8) (2021) 1971–1989.

[10] M. Levin, C. J. Martyniuk, The bioelectric code: An ancient computational medium for dynamic control of growth and form, Biosystems 164 (2018) 76–93.

[11] M. H. Baslow, The languages of neurons: an analysis of coding mechanisms by which neurons communicate, learn and store information, Entropy 11 (4) (2009) 782–797.

[12] D. S. Andres, The language of neurons: theory and applications of a quantitative analysis of the neural code, International Journal of Medical and Biological Frontiers 21 (2) (2015) 133.

[13] J. A. Pruszynski, J. Zylberberg, The language of the brain: real-world neural population codes, Current opinion in neurobiology 58 (2019) 30–36.

[14] R. Eckert, Y. Naitoh, K. Friedman, Sensory mechanisms in paramecium. i, J. exp. Biol 56 (1972) 683–694.

[15] M. Bingley, Membrane potentials in amoeba proteus, Journal of Experimental Biology 45 (2) (1966) 251–267.

[16] S. Ooyama, T. Shibata, Hierarchical organization of noise generates spontaneous signal in paramecium cell, Journal of theoretical biology 283 (1) (2011) 1–9.

[17] A. Hanson, Spontaneous electrical low-frequency oscillations: a possible role in hydra and all living systems, Philosophical Transactions of the Royal Society B 376 (1820) (2021) 20190763.

[18] T. Iwamura, Correlations between protoplasmic streaming and bioelectric potential of a slime mold, Physarum polycephalum, Shokubutsugaku Zasshi 62 (735-736) (1949) 126–131.

[19] N. Kamiya, S. Abe, Bioelectric phenomena in the myxomycete plasmodium and their relation to protoplasmic flow, Journal of Colloid Science 5 (2) (1950) 149–163.

[20] K. Trebacz, H. Dziubinska, E. Krol, Electrical signals in long-distance communication in plants, in: Communication in plants, Springer, 2006, pp. 277–290.

[21] J. Fromm, S. Lautner, Electrical signals and their physiological significance in plants, Plant, cell & environment 30 (3) (2007) 249–257.

[22] M. R. Zimmermann, A. Mithöfer, Electrical long-distance signaling in plants, in: Long-Distance Systemic Signaling and Communication in Plants, Springer, 2013, pp. 291–308.

[23] C. L. Slayman, W. S. Long, D. Gradmann, “action potentials” in neu-rospora crassa, a mycelial fungus, Biochimica et Biophysica Acta (BBA)-Biomembranes 426 (4) (1976) 732–744.

[24] S. Olsson, B. Hansson, Action potential-like activity found in fungal mycelia is sensitive to stimulation, Naturwissenschaften 82 (1) (1995) 30–31.

[25] A. Adamatzky, On spiking behaviour of oyster fungi pleurotus djamor, Scientific reports 8 (1) (2018) 1–7.

[26] A. Adamatzky, A. Gandia, On electrical spiking of ganoderma resinaceum, Biophysical Reviews and Letters 0 (0) (0) 1–9. arXiv:https://doi.org/10.1142/S1793048021500089, doi:10.1142/S1793048021500089. URL https://doi.org/10.1142/S1793048021500089

[27] A. Adamatzky, Language of fungi derived from their electrical spiking activity, Royal Society Open Science 9 (4) (2022) 211926.

[28] M. M. Dehshibi, A. Adamatzky, Electrical activity of fungi: Spikes detection and complexity analysis, Biosystems 203 (2021) 104373.

[29] J. Dons, O. De Vries, J. Wessels, Characterization of the genome of the basidiomycete schizophyllum commune, Biochimica et Biophysica Acta (BBA)-Nucleic Acids and Protein Synthesis 563 (1) (1979) 100–112.

[30] R. FitzHugh, Impulses and physiological states in theoretical models of nerve membrane, Biophysical journal 1 (6) (1961) 445–466.

[31] J. Nagumo, S. Arimoto, S. Yoshizawa, An active pulse transmission line simulating nerve axon, Proceedings of the IRE 50 (10) (1962) 2061–2070.

[32] A. M. Pertsov, J. M. Davidenko, R. Salomonsz, W. T. Baxter, J. Jalife, Spiral waves of excitation underlie reentrant activity in isolated cardiac muscle., Circulation research 72 (3) (1993) 631–650.

[33] G. W. Beeler, H. Reuter, Reconstruction of the action potential of ventricular myocardial fibres, The Journal of physiology 268 (1) (1977) 177–210.

[34] M. Tlalka, S. Watkinson, P. Darrah, M. Fricker, Continuous imaging of amino-acid translocation in intact mycelia of phanerochaete velutina reveals rapid, pulsatile fluxes, New Phytologist 153 (1) (2002) 173–184.

[35] L. F. Jaffe, Fast calcium waves, Cell calcium 48 (2-3) (2010) 102–113.

[36] R. Lopez-Franco, S. Bartnicki-Garcia, C. E. Bracker, Pulsed growth of fungal hyphal tips., Proceedings of the National Academy of Sciences 91 (25) (1994) 12228–12232.

[37] S. L. Jackson, Do hyphae pulse as they grow?, The New Phytologist 151 (3) (2001) 556–560.

